# Escherichia coli K12 exhibits a ~50% longer lag phase, but no difference in log phase growth rate, under hypomagnetic conditions (19 nT)

**DOI:** 10.64898/2026.04.13.717819

**Authors:** Michael Montague, Alessandro Lodesani, Clarice D. Aiello

## Abstract

Previous investigations have explored the effects of hypermagnetic fields, that is, fields in excess of the Earth’s background geomagnetic field strength of approximately 50 *µ*T, on *Escherichia coli* (*E. coli*). Conversely, this study investigates the effects of hypomagnetic field conditions, that is, fields below the geomagnetic background intensity, on the growth of *E. coli* K12 by using a hypomagnetic chamber to shield cultures, with a measured residual magnetic field inside the chamber of 19 nT. When grown in rich media from a semi-anaerobic, stationary-phase starting culture under geomagnetic and hypomagnetic conditions, the lag phases of *E. coli* were approximately 86 minutes and 132 minutes, respectively. Despite this increase in lag phase, exceeding two *E. coli* doubling times, the log-phase growth rate of *E. coli* was identical under both geomagnetic and hypomagnetic conditions. In addition to demonstrating a biologically relevant sensitivity to magnetic field parameters in the hypomagnetic direction, this represents a much greater absolute magnetosensitivity, with a deviation of only 50 *µ*T between the hypomagnetic and geomagnetic conditions, than has previously been demonstrated for *E. coli*.

## 1. Introduction

In biological assays, the effects of magnetic fields on living systems are rarely well controlled. One organism that has been investigated repeatedly with respect to magnetic fields is *Escherichia coli* (*E. coli*) [1–9]. However, all of these studies have used either relatively strong static magnetic fields [1, 8], oscillating magnetic fields [9], or both [3, 6]. Even studies that examine the effects of comparatively weak magnetic fields on *E. coli* have applied fields orders of magnitude greater than the Earth’s geomagnetic field, which is approximately 50 *µ*T. For example, Binhi et al. [2] tested the effects of applied static magnetic fields ranging from ‘0 mT’ to 110 mT on *E. coli* K12. At the highest field strength examined, the cells were exposed to a magnetic field approximately 2000 times stronger than the geomagnetic field. However, even at the lowest nominally applied field of ‘0 mT’, the actual exposure still corresponded to the ambient geomagnetic background of about 50 *µ*T. The absence of prior hypomagnetic studies in *E. coli* likely reflects the fact that generating a magnetic field is relatively straightforward, requiring only a wire coil or a Halbach array of permanent magnets, whereas shielding a sample from ambient magnetic fields requires a hypomagnetic chamber constructed from mu-metal.

A systematic survey of the effects of magnetic fields on biological systems involves many dimensions of complexity, including organism, magnetic field strength and frequency, other growth conditions, and the duration of altered field exposure, to name only a few. The space of all possible combinations of these variables is too large to be searched practically using empirical laboratory methods, even with automation. Consequently, it is advantageous to identify natural points of reduced complexity within this space. One such point is a static field with an absolute magnitude near 0 T that is maintained for the entire duration of the experiment, in contrast to the default geomagnetic field of approximately 50 *µ*T. These are referred to as ‘hypomagnetic’ conditions and do not require an arbitrary choice of field strength or frequency. Similarly, by comparing organisms maintained either in hypomagnetic conditions or in ambient geomagnetic conditions for the full duration of the experiment, there is no need to choose an arbitrary exposure time.

Rather than applying an additional field on top of the geomagnetic field, a baseline magnetic sensitivity should first be established under the most minimal and least arbitrary condition: a field strength near 0 T. For this reason, we investigate the effects of hypomagnetic field conditions, that is, field conditions with an absolute intensity below that of the geomagnetic field, on a model organism. Previous experiments demonstrated that hypomagnetic conditions alter the development of *Xenopus laevis* [10]. The study presented here shows, using *E. coli*, that sensitivity to hypomagnetic conditions is not limited to *Xenopus*. This work is intended as a starting point for future studies aimed at resolving the biological consequences of hypomagnetic conditions using *E. coli* as a model organism and higher resolution methods such as transcriptomics.

## 2. Materials and Methods

### 2.1. Strain and culture conditions

*E. coli* (Migula) Castellani and Chalmers 29425 K12 was obtained from ATCC [11]. Colonies were isolated on LB agar plates and grown in unsupplemented liquid LB medium (Lennox formulation; pH = 7.0), prepared according to the manufacturer’s instructions [12]. All cultures were incubated at 37 °C under ambient CO_2_ concentrations and agitated at 100 cycles per minute. Liquid cultures were grown in 25 mL volumes in 50 mL opaque (black) conical tubes held fully horizontally, such that all experiments were conducted under very low light exposure.

### 2.2. Growth curve measurements

Starting cultures of *E. coli* were established 24 hours prior to the growth curve experiments and allowed to reach stationary phase under geomagnetic conditions in sealed 50 mL conical tubes. Under these conditions, the starting culture naturally reached an optical density at 600 nm (OD_600_) of approximately 1.4 as oxygen in the sealed tube was depleted. As in the main time-course experiment, cultures were grown in 25 mL of LB in horizontal 50 mL opaque (black) conical tubes at 37 °C. On the day of the experiment, 10 mL of the starting culture was mixed with 200 mL of fresh LB and divided into eight 25 mL aliquots in separate conical tubes. Four tubes were placed under hypomagnetic incubation conditions at 37 °C, and four were placed under geomagnetic incubation conditions at 37 °C. Optical density at 600 nm (OD_600_) measurements were taken at 0, 30, 60, 90, 150, 210, 270, 330, and 390 minutes by removing 1 mL of medium and measuring it with an Eppendorf BioPhotometer. Growth rates were determined by identifying the regions of the average growth curves for the hypomagnetic and geomagnetic cultures with the maximum slopes (150–210 minutes for the geomagnetic cultures and 210–270 minutes for the hypomagnetic cultures) and calculating those slopes.

Care was taken to keep samples outside the incubator and hypomagnetic chambers for as little time as possible and to keep tubes open during sampling, and thus exposed to light, for as brief a period as possible. The average time outside was on the order of 90 seconds for all samples at each time point.

### 2.3. Hypomagnetic field environment and field measurements

A Twinleaf MS-1L hypomagnetic chamber installed inside an agitating incubator provided the hypomagnetic conditions used to test *E. coli*. Because electronic devices used to maintain environmental controls within the chamber would generate their own magnetic fields, it was more practical to place the hypomagnetic chamber inside an incubator than to place an incubator inside a hypomagnetic chamber. The chamber was degaussed once and measured to have an internal magnetic field strength of 19.9 nT. No active field compensation was used in these experiments. The residual field strength was measured with a QuSpin sensor using QuSpin 2FM UI V7.6.2 software.

## 3. Results

The growth curves of *E. coli* K12 exhibited identical log-phase growth rates of 0.013 OD_600_ per minute, but the lag phase of the hypomagnetic cultures was substantially longer. The observed lag phase under geomagnetic conditions was 86.6 minutes, consistent with the use of a stationary-phase starting culture [13], whereas the lag phase of otherwise identical cultures grown under hypomagnetic conditions was 132.2 minutes. This represents a 52.6% increase in lag phase and exceeds two doubling times of *E. coli* during log-phase growth under these conditions, as shown in Fig. 1.

**Fig. 1:**
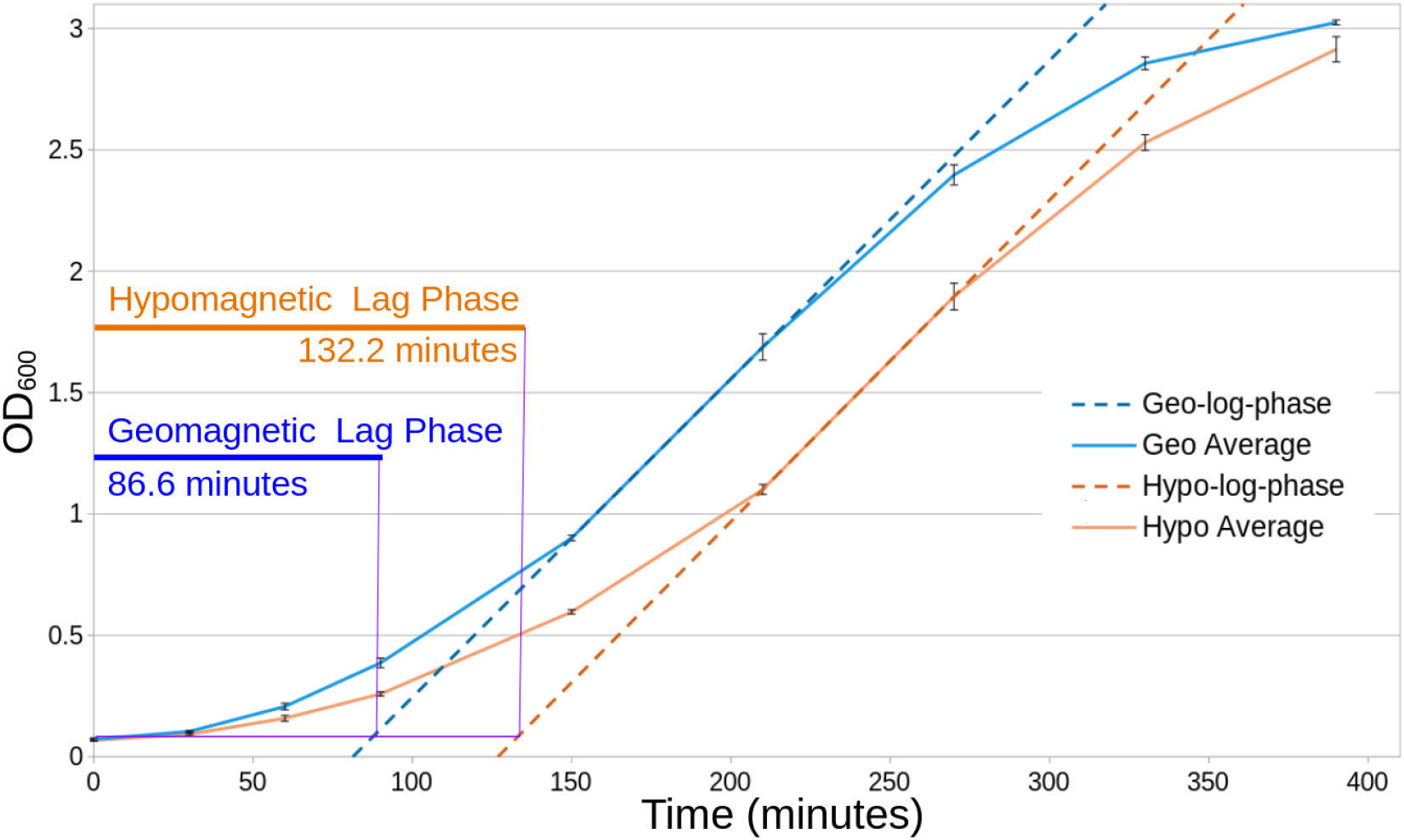
Growth curves of *E. coli* K12 under geomagnetic and hypomagnetic conditions show increased lag-phase for bacteria under hypomagnetic conditions. Hypomagnetic cultures exhibit a longer lag phase (132.2 vs. 86.6 minutes). Log-phase growth rates are identical, at 0.013 OD_600_ per minute.

A total of eight tubes per condition were analyzed across two independent experiments conducted on separate days using separate batches of media, thereby providing both technical replicates (shared starting culture and media batch within an experiment) and biological replicates (different starting cultures and media batches between experiments).

## 4. Discussion

### These results are significant for several independent reasons

First, the difference between the geomagnetic and hypomagnetic field strengths was only 50 *µ*T. This demonstrates a degree of absolute magnetosensitivity in *E. coli* that has not previously been reported. Previous studies have consistently focused on applied field strength differences in the mT range or greater [1–9]. Effects of weak static magnetic fields are predicted by the radical pair mechanism, one of the leading candidate mechanisms proposed to explain biological magnetosensitivity [14, 15]. Furthermore, this sensitivity falls broadly within the range of natural variation in the intensity of the Earth’s magnetic field at the surface, which, although averaging about 50 *µ*T, varies by location from approximately 20 *µ*T–65 *µ*T [16, 17].

Second, the hypomagnetic field condition represents a novel direction for probing magnetic field effects in *E. coli*. In considering the significance of a hypomagnetic, rather than hypermagnetic, effect on a living system, it is worth noting that the Earth’s surface is among the most magnetically exposed regions of near space. Consequently, all locations currently under consideration for crewed space expeditions exist naturally under what would be considered hypomagnetic conditions: the Moon (*<* 10 nT) [18], Mars (*<* 220 nT) [19], and low Earth orbit, where field strengths are on the order of tens of *µ*T but remain lower than those at the Earth’s surface and vary with orbital parameters [20].

Third, of the nine hypermagnetic field experiments mentioned above, six [1, 5–9] examined how *E. coli* growth kinetics or viability were affected by exposure to hypermagnetic fields, and all six reported reduced growth or viability. Although direct comparison is difficult due to differences in experimental conditions, the consistent observation of reduced growth or viability under hypermagnetic conditions, together with the prolonged lag phase we observed under hypomagnetic conditions, suggests that deviations from geomagnetic field strength in either direction are detrimental to *E. coli* growth, even if the underlying mechanisms are not the same.

Fourth, taken together with the identical log-phase growth rates of the geomagnetic and hypomagnetic cultures, the prolonged lag phase under hypomagnetic conditions suggests that these conditions do not affect the underlying viability of *E. coli*. Rather, the extended lag phase may result from interference with regulatory or signaling processes. This, in turn, suggests that magnetosensitivity in *E. coli*, at least in the hypomagnetic regime, may be primarily informational in nature. In other words, magnetic field sensing may contribute to the acquisition of environmental information that helps regulate biological states and processes.

Future directions for this research include characterizing the boundaries of the hypomagnetic effect on *E. coli*. In part, this will involve the use of higher-resolution approaches, such as multi-omics, to determine which genes are differentially expressed. Although the absolute sensitivity demonstrated here is much greater than previously expected, it should be examined further with respect to both magnetic field strength and the duration of hypomagnetic exposure. The mechanism underlying magnetosensitivity may also be investigated by altering the conditions of hypomagnetic exposure. For example, although this experiment was performed under dark conditions, some theories of biological magnetosensitivity, such as the radical pair mechanism [14], suggest that light may play a role. Determining whether the presence or absence of light alters magnetosensitivity may therefore be informative. Other potential confounding factors, such as aerobic versus anaerobic conditions, may also help clarify the role of reactive oxygen species (ROS). Finally, although *E. coli* is a well-studied Gram-negative bacterium, it represents only a small portion of the genetic diversity of life on Earth. Similar hypomagnetic studies should therefore be carried out in other microbial model organisms, such as *B. subtilis* (Gram-positive bacterium), *H. volcanii* (archaeon), and *S. cerevisiae* (eukaryote), with the eventual goal of constructing a survey of magnetosensitivity across the phylogenetic diversity of life.

## 5. Author Contributions

Following the CRediT taxonomy [21], the authors confirm the following contributions:

1. Michael Montague: conceptualization; investigation; data curation; methodology; validation; formal analysis; writing (original draft); and visualization
2. Alessandro Lodesani: conceptualization; methodology; investigation; and writing (review and editing)
3. Clarice D. Aiello: conceptualization; methodology; writing (review and editing); supervision; project administration; and funding acquisition

## 6. Acknowledgements

We would like to thank Morgan L. Sosa for her assistance with formatting, revising, and editing the manuscript.

This project was funded by a grant from the [Quantum Biology DAO](https://www.quantumbiology.xyz/).

The [Quantum Biology Institute](https://www.quantumbiology.org/) is a California nonprofit (501(c)(3)) focused research organization [22] that performs basic research underpinning the quantum biology field in an open-science fashion. It is part of the [Quantum Biology Ecosystem](https://www.quantumbiology.eco/).

